# Early electrical stimulation promotes functional recovery after volumetric muscle loss

**DOI:** 10.64898/2026.03.01.708642

**Authors:** Samuel Gershanok, Anne Behre, Rifeng Jin, Sofya Vinokurova, John Blount, Raghav Garg, Alpaslan Ersöz, Liyang Wang, Seonghan Jo, Daniel Ranke, Mangesh Kulkarni, Devora Cohen-Karni, Adam Feinberg, Douglas Weber, Bryan Brown, Tzahi Cohen-Karni

## Abstract

Volumetric muscle loss (VML) injuries overwhelm the inherent regenerative capacity of skeletal muscle, causing persistent functional deficits with no routinely effective therapies. Electrical stimulation (ES) has been shown to preserve muscle structure in other injury models, but technical barriers have prevented daily delivery during the acute post-injury window when critical regenerative programs are established. Here, we developed a fully implantable bioelectronic system with nanoporous platinum-modified electrodes enabling daily therapeutic stimulation and electromyographic recording without repeated anesthesia in a rat tibialis anterior VML model. Animals receiving ES during the acute post-injury period (10 sessions over days 0–14) showed sustained functional improvement, reaching 90% of baseline torque at 8 weeks compared to 71% in unstimulated controls. This recovery reflected enhanced remodeling of injured muscle rather than synergistic muscle compensation. Histological analysis revealed coordinated early increases in vascularization, pro-regenerative macrophages, and satellite cells. These findings establish early ES as a promising intervention for promoting muscle regeneration after catastrophic injury.

## Introduction

Volumetric muscle loss (VML) involves irrecoverable loss of muscle tissue that exceeds the body’s endogenous regenerative capacity.^1^ These injuries are characterized by chronic inflammation, fibrotic scarring and impaired neuromuscular repair, resulting in persistent functional deficits.^2–5^ VML represents a substantial clinical burden: muscle conditions account for 65% of disability ratings in military personnel with orthopedic trauma and VML occurs commonly in civilian populations secondary to open fractures, gunshot wounds, and surgical resections.^4, 6^ Current clinical interventions, including physical rehabilitation,^7,8^ orthotic bracing,^9,10^ and autologous muscle flaps^1,11^ remain largely palliative because none of these approaches reliably restores lost muscle architecture and force-generating function. There is therefore a pressing need for therapies that actively promote coordinated neuromuscular regeneration rather than simply mitigate disability.

Electrical stimulation (ES) delivers controlled electrical pulses through surface or implanted electrodes to depolarize peripheral nerves and/or muscle fibers, eliciting repeated contractions and activity-dependent signaling that can shape muscle performance and remodeling.^12–14^ In smaller, recoverable neuromuscular injuries, such as contusions, strains, and nerve crush injuries that would eventually heal without intervention, daily ES initiated during the acute post-injury interval (0–2 weeks) accelerates the restoration of muscle structure and function and supports neuronal integrity.^15–18^ and supports neuronal integrity.^19–21^ Whether early ES can redirect the dysfunctional healing pattern characteristic of VML, however, has not been established.

The rationale for targeting this early window lies in the convergence of immune, myogenic, and vascular programs that together determine long-term regenerative outcomes. During acute post-injury healing, macrophage polarization from pro-inflammatory to pro-regenerative states, satellite (muscle progenitor) cell activation and expansion, and microvascular remodeling occur in parallel and are each sensitive to electrical and mechanical cues.^22–26^ Preclinical evidence suggests that appropriately timed ES can accelerate macrophage phenotype switching and enhance satellite cell proliferation in non-VML injury contexts, but whether these effects translate to the VML setting is unknown.

Realizing this potential in VML, however, has been hindered by two practical barriers. First, most rodent VML studies deliver ES using transcutaneous or percutaneous needle electrodes that typically require repeated general anesthesia, constraining dosing frequency and preventing sustained intervention during the acute inflammatory phase.^2, 27–29^ Needle-based approaches necessitate anesthesia because electrode insertion and precise positioning require the animal to remain immobile, and repeated percutaneous procedures without anesthesia would cause unacceptable pain and stress. Second, functional readouts can be confounded by hypertrophic compensation of intact synergistic muscles, complicating attribution of torque improvements to regeneration within the injured muscle.^2, 29^ Resolving both limitations is essential to determine whether early, daily suprathreshold ES (stimulation strong enough to reliably activate nerve and muscle) can meaningfully improve neuromuscular recovery after VML.

Here, we present a fully implantable bioelectronic approach that overcomes both barriers by enabling repeated stimulation without anesthesia and longitudinal electrophysiological monitoring in a rat tibialis anterior VML model. Through paired functional, electromyographic (EMG), and histological assessments over 8 weeks, we show that immediate post-injury ES enhances neuromuscular recovery and is associated with favorable shifts in macrophage polarization, satellite cell activity, and vascular remodeling, supporting early stimulation as a therapeutic strategy for VML.

## Results

### Implanted VML Bioelectronic System

To test whether electrical stimulation (ES) delivered early after injury can improve muscle recovery, we developed a fully implantable system for rats that delivers precisely timed electrical pulses and records muscle activity without requiring repeated anesthesia **(Fig. 1a)**. We used a well-established volumetric muscle loss (VML) model in which a defined portion 20% the tibialis anterior (TA) muscle is surgically removed **(Fig. S1)**.^2^ Immediately after injury, we implanted small, flexible electrodes directly onto the remaining muscle adjacent to the defect site. Because these electrodes remain in place throughout recovery, the system can deliver stimulation strong enough to reliably activate nerve and muscle tissue, following protocols previously shown to promote healing in other neuromuscular injury models.^16, 17, 20, 21, 30^ This represents a key advance over prior VML studies using temporary needle electrodes, which require animals to be anesthetized for each treatment session.^27–29^

**Figure 1.**
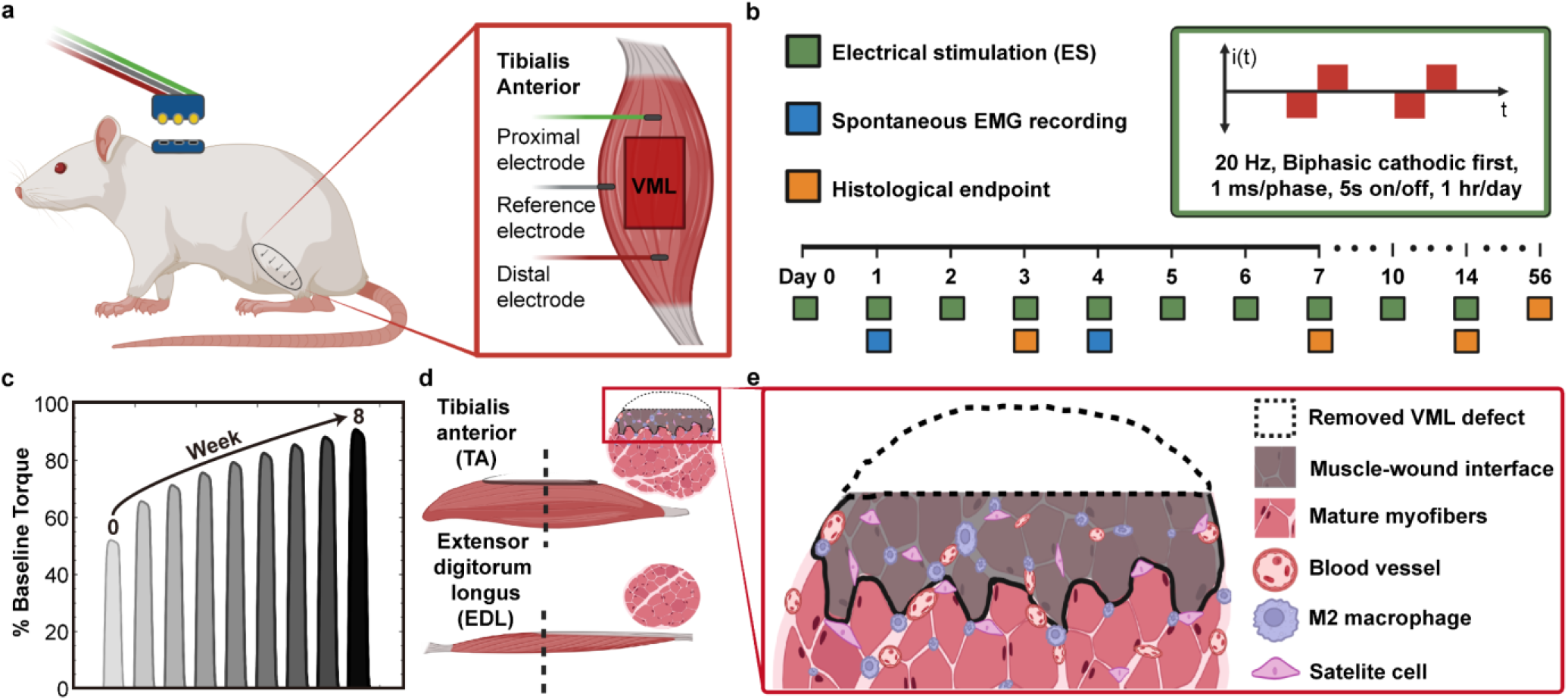
Fully implantable system for electrical stimulation after volumetric muscle loss. **(a)** Schematic of the rat tibialis anterior (TA) VML model with electrode placement for stimulation and EMG recording. **(b)** Stimulation parameters and timeline for ES delivery, EMG acquisition, and histological endpoints. **(c)** Weekly maximum isometric dorsiflexion torque as a measure of functional recovery. **(d)** Myofiber cross-sectional area (CSA) in the injured TA and extensor digitorum longus (EDL) to assess compensatory hypertrophy. **(e)** Representative Masson’s trichrome and immunohistochemical staining at the muscle–wound interface showing ES-associated changes in vascular density, M2 macrophage abundance, and satellite cell activation.

Using this system, we delivered one-hour stimulation sessions during the acute post-injury period (days 0–7, 10, and 14; 10 sessions total; **Fig. 1b**). We studied three groups: VML with stimulation (VML+ES, n=28), VML without stimulation (VML+NS, n=28), and uninjured controls receiving stimulation (ND+ES, n=8). Male and female rats were distributed across all groups in approximately equal numbers. We tracked functional recovery through weekly torque measurements **(Fig. 1c)**, assessed muscle fiber size in both the injured TA and the extensor digitorum longus (EDL), an adjacent synergist that contributes to dorsiflexion and can hypertrophy to compensate for TA loss **(Fig. 1d)**, and performed immunohistochemistry at multiple timepoints to examine inflammation, blood vessel formation, and muscle stem cell activity **(Fig. 1e)**.

### Electrode characterization and in vivo stability

We modified electrodes with nanoporous platinum (NanoPt) to ensure safe, stable stimulation. Initial testing with unmodified electrodes in muscle-mimicking tissue produced gas bubbles and surface oxidation **(Fig. S2a)**, electrochemical byproducts that can damage both electrodes and surrounding tissue.^31–33^ In contrast, NanoPt-modified electrodes showed no gas evolution or surface degradation under identical conditions **(Fig. S2b)**. Scanning electron microscopy confirmed that NanoPt modification preserves the flexible multistrand architecture **(Fig. 2a.i)** while introducing a highly porous surface texture **(Fig. 2a.ii)**. This increased surface area lowers electrical resistance at the electrode–tissue interface and allows higher currents to be delivered safely **(Fig. S3)**. Importantly, NanoPt-modified electrodes subjected to 48 hours of continuous stimulation in phantom tissue showed no change in charge-storage capacity **(Fig. 2b, S2c–d)**, indicating electrochemical stability suitable for chronic in vivo use.

**Figure 2:**
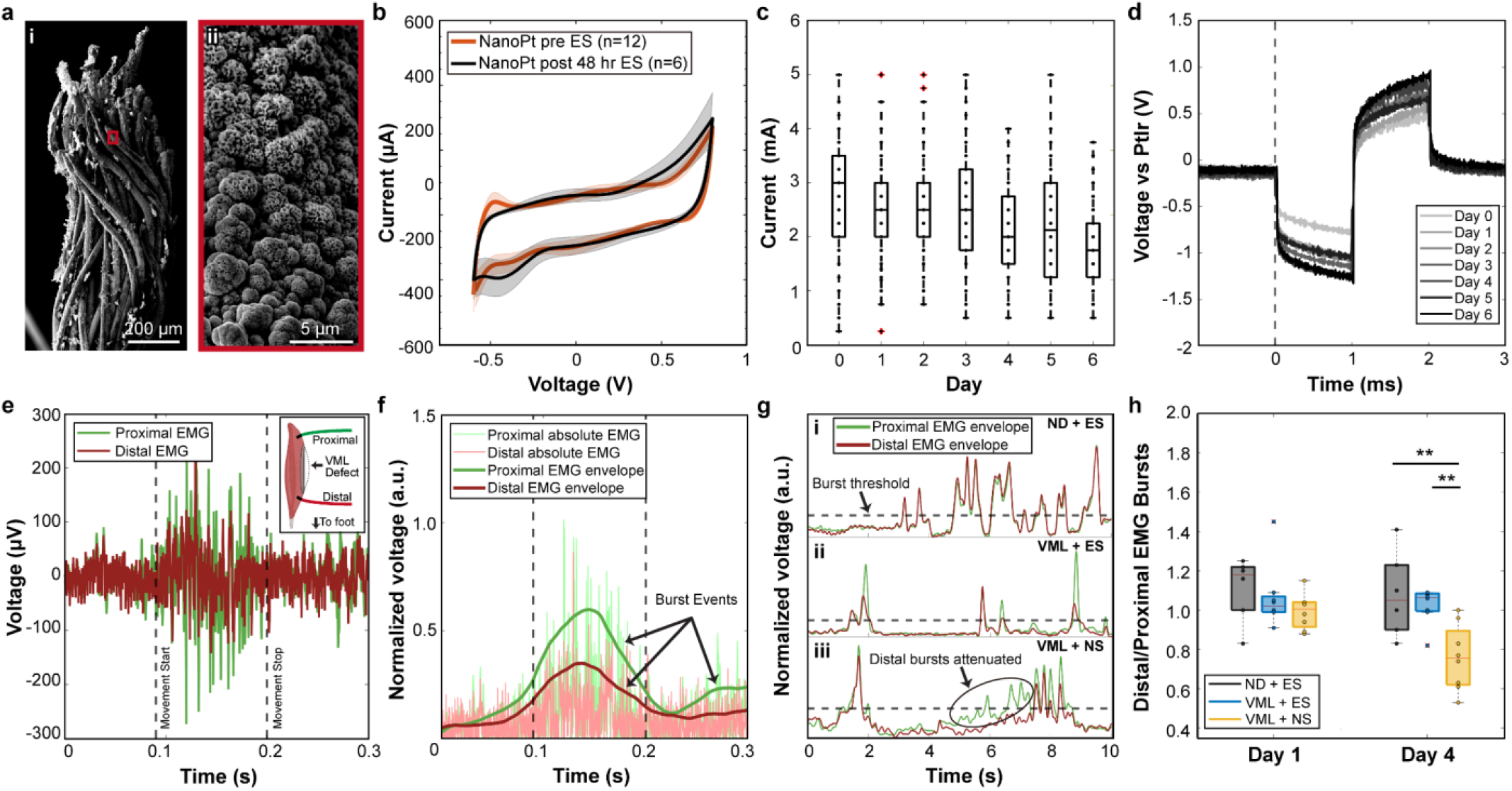
NanoPt-modified electrodes provide stable stimulation and reveal early protective effects of ES on neuromuscular excitability. **(a)** Scanning electron micrographs showing that NanoPt modification **(i)** preserves multistrand electrode architecture while **(ii)** introducing a porous nanoscale surface. **(b)** Cyclic voltammetry of NanoPt-modified electrodes in phantom tissue before and after 48 h of continuous stimulation, demonstrating stable charge-storage capacity. **(c)** Current required to elicit visible muscle contraction across the seven-day stimulation protocol (n = 10–16). **(d)** Voltage transients recorded at maximal current relative to the implanted reference electrode from day 0 through day 6 (n = 5–10). **(e)** Raw EMG traces recorded during spontaneous locomotion from electrodes flanking the VML defect **(inset). (f)** Processed EMG burst envelopes with quantified burst events. (g) Representative burst envelopes showing distal and proximal firing dynamics at day 4 across **(i)** ND+ES, **(ii)** VML+ES, and **(iii)** VML+NS groups. **(h)** Distal-to-proximal EMG burst ratio at days 1 and 4 post-injury (n = 6 ND+ES; n = 8 VML+ES and VML+NS). Voltage transients show mean ± s.d. (shaded). Box plots display median, interquartile range, and min–max; outliers marked by red “+”. \*\**P<* 0.01, two-tailed Student’s t-test.

To confirm that stimulation reliably activated muscle throughout the treatment window, we recorded current thresholds during each session. Tetanic contractions were evoked by passing current between the distal and proximal electrodes, with amplitude adjusted every 10 minutes to maintain visible contraction **(Fig. 2c)**. Within individual sessions, current typically had to be increased over time, consistent with established short-term reductions in neuromuscular excitability during repeated activation.^34, 35^ Importantly, the current required to evoke contraction at the start of each session did not differ between days 0 and 6, indicating that nerve and muscle excitability fully recovered between sessions without cumulative fatigue.^36, 37^

To assess whether the electrode–tissue interface remained stable over repeated use, we analyzed voltage waveforms recorded during stimulation **(Fig. 2d)**. Throughout the seven-day protocol, waveforms remained charge-balanced (equal positive and negative phases, preventing tissue damage) with no drift in the resistive or capacitive components of the signal. Such drift would indicate electrode surface degradation or unstable positioning, neither of which was observed.^38–40^ These stable voltage profiles, combined with consistent current thresholds across days, confirm that the implanted electrodes maintained reliable electrical contact with the tissue throughout the acute treatment period.

### Spontaneous EMG reveals early protective effects of ES

Having confirmed electrode stability, we next asked whether the implanted system could detect ES-induced changes in neuromuscular function. A key advantage of our implantable platform is the ability to record spontaneous EMG activity during natural, voluntary movement rather than evoked responses under anesthesia, providing a more physiologically relevant measure of functional recovery. To investigate how ES affects acute neuromuscular function, we recorded spontaneous EMG activity on days 1 and 4 post-injury. EMG burst activity reflects voluntary muscle activation during natural movement^41, 42^ and is attenuated in rodents with TA injury.^43^ We recorded activity from electrodes positioned proximally and distally relative to the VML defect as animals moved freely in their cages **(Fig. 2e)**. From these recordings, we generated burst envelopes to quantify the number of burst events detected at each electrode location **(Fig. 2f–g, S4a)**, as well as burst duration **(Fig. S4b)** and amplitude **(Fig. S4c)**. At day 1, burst characteristics were identical across all three groups. By day 4, however, VML+NS animals showed significantly fewer burst events at the distal electrode compared to the proximal electrode, a reduction not observed in either ND+ES (*P =* 0.0087) or VML+ES (*P =* 0.0070) groups **(Fig. 2h)**.

This pattern reflects the anatomy of intramuscular innervation. The TA receives motor axons from the deep peroneal nerve, which enter proximally and branch extensively throughout the muscle.^44^ Because our mid-belly VML defect unavoidably severs some of these intramuscular branches, motor units distal to the injury lose their neural input. At day 1, distal nerve terminals and muscle fibers remain transiently excitable because Wallerian degeneration, the process by which severed axons distal to the site of injury break down, takes 24–48 hours to complete.^45, 46^ Without intervention, this degeneration leads to muscle fiber atrophy,^47, 48^ explaining the selective loss of distal EMG activity in VML+NS animals by day 4. In contrast, ES-treated animals maintained distal burst activity, suggesting preserved excitability of both nerve and muscle tissue across the defect. This protective effect aligns with prior evidence that electrical stimulation supports axonal integrity, enhances neurotrophic signaling, and maintains Schwann cell organization.^49, 50^ Importantly, these differences emerged during volitional activity, suggesting that ES preserves not only the capacity for muscle activation but also its functional integration into natural motor behavior. These findings suggest that early ES may counteract denervation-induced deterioration by sustaining the excitability of tissue distal to the injury site.

### Early ES improves functional recovery after VML

To determine whether early ES improves long-term function, we measured maximum isometric dorsiflexion torque weekly through week 8 **(Fig. 3a, S5)**. Torque was normalized to each animal’s pre-injury baseline to account for individual variation. Immediately after VML, all injured animals showed similar deficits (~40% of baseline; **Fig. 3b–c**). By week 2, VML+ES animals had recovered significantly more torque than VML+NS controls (+32%, *P =* 0.0004), and this advantage persisted through week 8 (+27%, *P =* 0.0080). VML+NS animals plateaued at ~70% of baseline, consistent with the persistent 28–33% deficit reported in this model.^2^ In contrast, VML+ES animals recovered to levels statistically indistinguishable from uninjured ND+ES controls by week 8 (*P =* 0.158).

**Figure 3:**
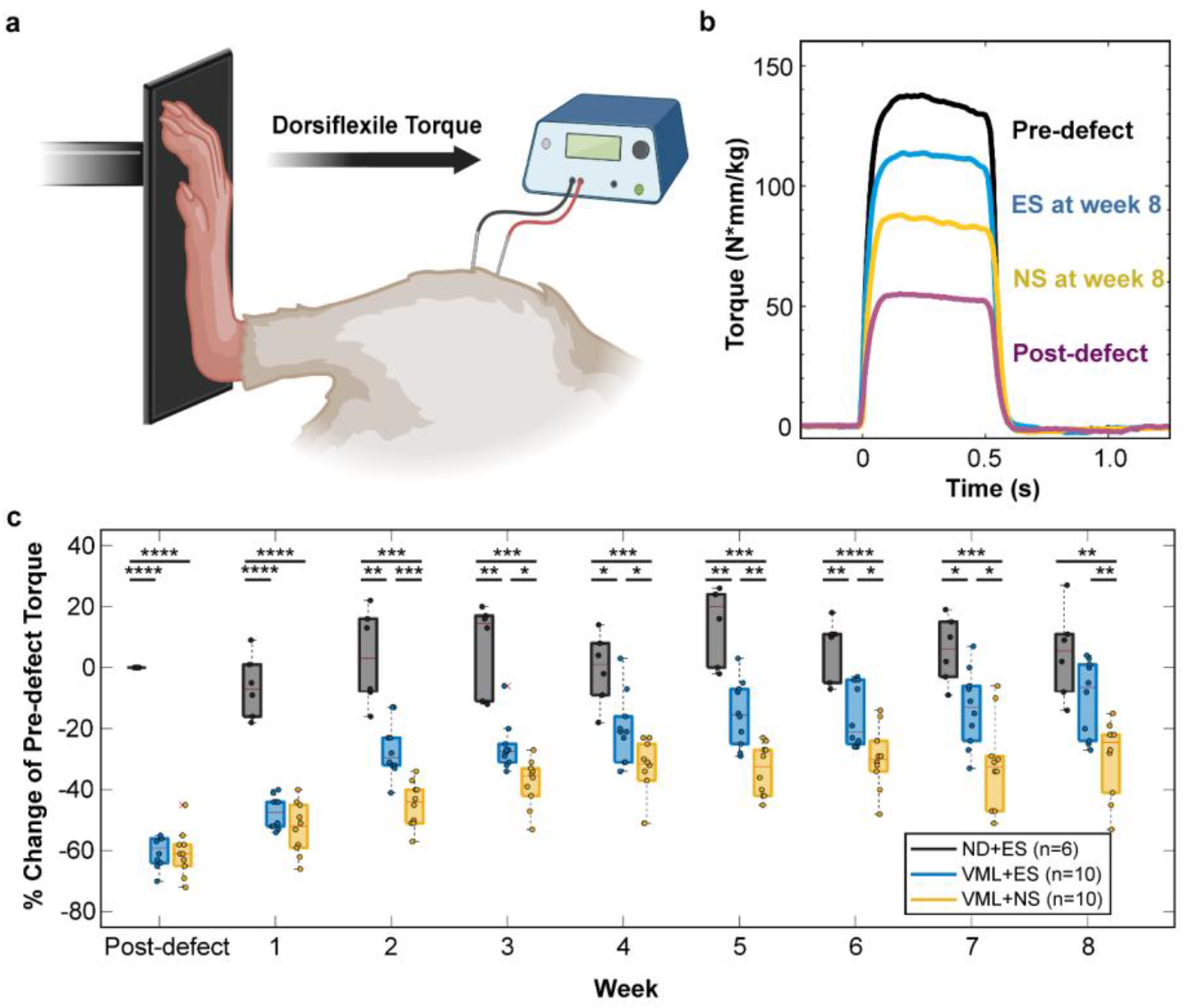
Early ES restores dorsiflexion torque after VML. **(a)** Schematic of torque measurement using a force gauge coupled to a footplate, with peroneal nerve stimulation via needle electrodes. **(b)** Representative torque traces at pre-defect, post-defect (day 0), and week 8 for ND+ES, VML+ES, and VML+NS groups. **(c)** Maximum tetanic torque normalized to pre-defect values across 8 weeks (n indicated in legend). Box plots show median, interquartile range, and min–max; outliers marked by red “×”. Two-way ANOVA with post-hoc t-tests. \**P<* 0.05, \*\**P<* 0.01, \*\*\**P<* 0.001, \*\*\*\**P<* 0.0001.

To confirm that these gains reflected injury-specific effects rather than ES-induced hypertrophy, we examined torque in uninjured animals receiving stimulation. ND+ES torque did not change across the eight-week protocol (one-way ANOVA, *P =* 0.207), indicating that ES alone does not augment force production in intact muscle. Together, these data demonstrate that early ES yields a sustained ~30% functional improvement, restoring dorsiflexion strength to near-normal levels following VML.

### ES preserves TA mass without inducing EDL compensation

To determine whether the functional gains observed with ES reflect true TA remodeling rather than synergist compensation, we measured wet muscle mass and myofiber cross-sectional area (CSA) of both the TA and EDL at week 8. The extensor digitorum longus (EDL), which lies adjacent to the TA and contributes to dorsiflexion, can hypertrophy after VML to partially compensate for lost TA tissue.^2^ Measuring both muscles therefore allows us to distinguish TA-specific regeneration from compensatory adaptation. For mass measurements, we compared three groups: uninjured animals receiving stimulation (ND+ES), VML with stimulation (VML+ES), and VML without stimulation (VML+NS). For CSA analysis, we compared VML groups against tissue from the contralateral (uninjured, unstimulated) limb (ND), which served as the baseline reference for normal fiber morphology.

ES prevented the TA mass loss characteristic of VML. Defect-to-contralateral TA mass ratios were significantly lower in VML+NS animals compared to both ND+ES (*P =* 0.0097) and VML+ES (*P =* 0.0157) groups, while VML+ES and ND+ES did not differ statistically (*P =* 0.172; **Fig. 4a.i**). This indicates that early stimulation preserved TA mass at near-normal levels.

**Figure 4:**
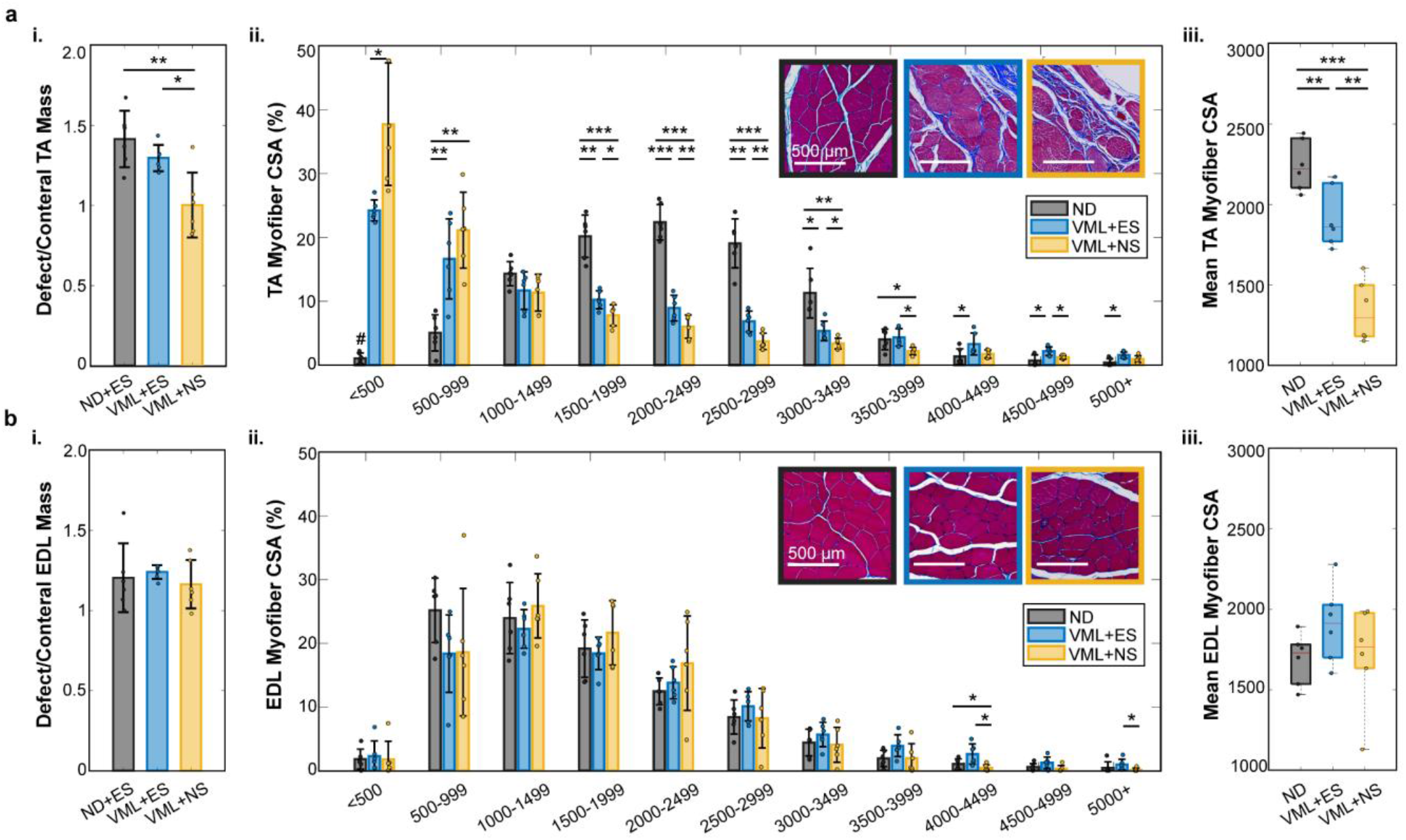
ES preserves TA mass and fiber size without differential EDL compensation. Quantitative analysis of **(a)** TA and **(b)** EDL at week 8. **(i)** Defect-to-contralateral wet mass ratio for ND+ES, VML+ES, and VML+NS groups. **(ii)** Myofiber CSA distribution compared to contralateral baseline (ND), with representative H&E images (insets; scale bar, 500 µm). **(iii)** Mean myofiber CSA. n = 6 biological replicates per group. Bar graphs show mean ± s.d. Box plots show median, interquartile range, and min–max. \**P<* 0.05, \*\**P<* 0.01, \*\*\**P<* 0.001, two-tailed t-test.

EDL mass increased approximately 20% above contralateral values across all groups, consistent with compensatory hypertrophy reported in this model.^2^ Importantly, EDL mass ratios did not differ among ND+ES, VML+ES, and VML+NS animals (one-way ANOVA, *P* = 0.69, **Fig. 4b.i**), indicating that ES-driven torque improvements cannot be attributed to differential synergist compensation.

Myofiber CSA analysis revealed that ES shifted the TA fiber size distribution toward larger fibers. Both VML groups showed increased proportions of small fibers (<1000 µm^2^) concentrated at the defect edge, reflecting localized regeneration at the wound boundary^51^ **(Fig. 4a.ii, insets)**. However, VML+ES animals exhibited a greater proportion of large fibers (>3500 µm^2^) and higher mean CSA compared to VML+NS (*P* = 0.0023; **Fig. 4a.iii**). In contrast, EDL fiber size distributions and mean CSA did not differ between VML+ES and VML+NS groups and remained similar to ND baseline values **(Fig. 4b.ii–iii)**, confirming that the synergistic muscle responded similarly regardless of treatment.

Together, these data demonstrate that ES preserves TA mass and promotes larger fiber sizes without differentially affecting the EDL. This supports the conclusion that the observed torque improvements reflect enhanced TA remodeling rather than synergistic compensation.

### ES promotes coordinated regenerative remodeling at the injury site

To examine the cellular changes associated with ES-driven functional improvement, we performed histological analysis. t multiple timepoints (pre-defect, day 3, weeks 1, 2, and 8). For each sample, we quantified tissue composition and cell populations within a standardized 1 × 3 mm region of interest encompassing the defect site, adjacent remodeling tissue, and neighboring healthy muscle (**Fig. 5a**; see Methods).

**Figure 5:**
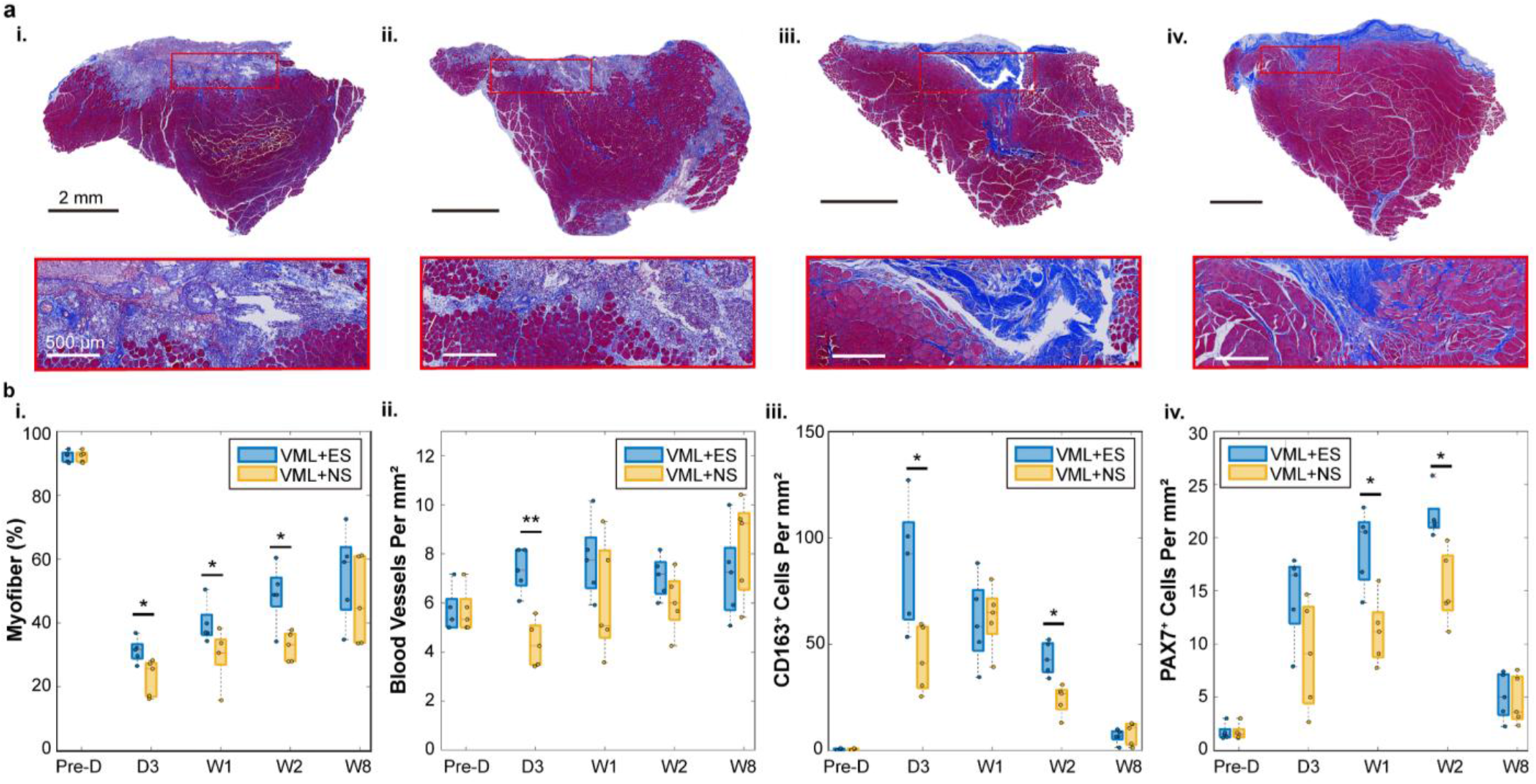
ES coordinates early regenerative remodeling after VML. **(a)** Representative Masson’s trichrome cross-sections at day 3 (**i**, VML+NS; **ii**, VML+ES) and week 8 (**iii**, VML+NS; **iv**, VML+ES). Outlined regions (1 × 3 mm) indicate the superficial TA area encompassing the defect site and adjacent tissue used for quantification; higher magnification shown below. **(b)** Quantification of **(i)** myofiber area percentage, **(ii)** blood vessel density (vessels >100 µm^2^), **(iii)** CD163^+^ (M2 macrophage) cell density, and **(iv)** Pax7^+^ (satellite cell) density across timepoints. n = 5 biological replicates per group, each derived from four technical replicates. Box plots show median, interquartile range, and min–max; outliers marked by red “×”. \**P<* 0.05, \*\**P<* 0.01, two-tailed t-test.

ES preserved myofiber content throughout the early post-injury period. Masson’s trichrome staining revealed that both VML groups lost myofiber area compared to pre-defect baseline, but VML+ES animals retained significantly more myofiber content than VML+NS at day 3 (*P =* 0.0441), week 1 (*P =* 0.0336), and week 2 (*P =* 0.0440; **Fig. 5b.i**).

ES enhanced early vascular remodeling. Quantification of blood vessels (>100 µm^2^ cross-sectional area, capturing arterioles and venules rather than capillaries) revealed significantly higher vessel density in VML+ES compared to VML+NS at day 3 (*P =* 0.0135; **Fig. 5b.ii**). This increase was driven primarily by young, developing vessels **(Fig. S6)**, consistent with prior reports of activity-induced vessel growth in skeletal muscle.^25, 52^

ES shifted macrophage populations toward a pro-regenerative phenotype. Following muscle injury, macrophages transition from a pro-inflammatory state that clears debris to a pro-regenerative (M2) state that supports tissue repair. Immunohistochemical quantification of CD163, a marker of M2 macrophages, showed significantly elevated CD163^+^ cell density in VML+ES versus VML+NS at day 3 (*P =* 0.0186) and week 2 (*P =* 0.0231; **Fig. 5b.iii, S7, S8**). This early shift toward pro-regenerative macrophages may help resolve inflammation and create a microenvironment conducive to muscle repair.

ES increased satellite cell abundance during the proliferative phase. Satellite cells are muscle-resident stem cells that normally lie dormant beneath muscle fibers but activate in response to injury, proliferating and fusing to repair damaged tissue. We quantified these cells using Pax7, a transcription factor expressed by quiescent and activated satellite cells. VML+ES animals displayed significantly more Pax7^+^ cells than VML+NS at week 1 (*P =* 0.0465) and week 2 (*P =* 0.0469; **Fig. 5b.iv, S9, S10**), indicating enhanced satellite cell activation or retention during the period when these cells contribute most actively to fiber regeneration.

Together, these findings suggest that early ES coordinates multiple regenerative programs during the critical acute-to-proliferative transition after VML, including preserved myofiber content, enhanced vascularization, M2 macrophage polarization, and expanded satellite cell populations.

## Discussion

This study demonstrates that early, daily electrical stimulation initiated after VML significantly improves long-term functional recovery. Using a fully implantable bioelectronic system that overcomes the repeated anesthesia requirement of prior approaches, we show that ES delivered during the acute post-injury period (10 sessions over two weeks) both accelerates functional recovery and restores dorsiflexion torque to near-normal levels by week 8. Critically, this functional benefit reflects enhanced TA remodeling rather than synergist compensation, as evidenced by preserved TA mass and fiber size without differential changes in the EDL. These findings address the gap identified at the outset: whether early ES can redirect the dysfunctional healing trajectory characteristic of VML.

The functional benefits of early ES appear to arise through preservation of neuromuscular structure during the acute post-injury window. ES increased myofiber content from day 3 through week 2 and maintained EMG burst propagation across the defect at day 4, suggesting that stimulation protects both muscle and nerve tissue from early degeneration. Because uninjured animals receiving ES (ND+ES) showed no torque gains, hypertrophy of pre-existing fibers was not the dominant mechanism. Instead, ES appears to preserve tissue integrity around the defect, leading to larger myofiber cross-sectional areas and fewer small, immature fibers at week 8. These observations support a model in which early ES enhances long-term recovery by protecting neuromuscular connectivity and accelerating functional restoration, thereby limiting secondary atrophy, rather than by inducing compensatory growth.

The cellular mechanisms likely involve coordinated effects on perfusion of the injury site and immune signaling. Electrical stimulation at the parameters used here (20 Hz, 1 ms pulse duration) has been shown to increase blood flow and vessel density in ischemic models.^53^ Our observation of increased developing vasculature at day 3 aligns with evidence that ES-induced electric fields activate vascular endothelial growth factor (VEGF) signaling in endothelial^54^ and muscle^55^ cells, promoting rapid vessel formation.^56^ While we did not directly measure VEGF expression, the temporal pattern of vessel development is consistent with this mechanism. In parallel, we observed elevated CD163^+^ M2 macrophages, a population that resolves inflammation, stimulates new vessel growth, and guides satellite cell differentiation following recoverable muscle injury.^22^ Prior work indicates that ES can directly bias macrophage polarization toward this pro-regenerative phenotype^57–59^ and that macrophages have both paracrine and physical interactions with satelite cells that support both fiber regeneration and vascularization.^60, 61^ In VML, where regeneration is often limited by poorly coordinated inflammation and inadequate vascular support,^62^ ES may help synchronize these programs during the critical early window.

Several limitations warrant consideration. Our partial-thickness VML model may not capture the full complexity of larger, multi-muscle, or multi-tissue injuries involving skin, bone, and neurovascular structures. Indeed, prior studies using biweekly ES starting 72 hours post-injury in a more severe full-thickness model showed no functional benefit,^28^ suggesting that timing, dosing, and injury severity interact in ways that require further investigation. Additionally, while we observed correlations between early cellular changes and later functional outcomes, direct causal links remain to be established. Future work should examine whether this early ES protocol can be translated to larger injuries and, ultimately, to clinical rehabilitation strategies for patients with traumatic muscle loss. Addressing these questions will clarify how early ES can be optimized as a foundational intervention for VML.

## Conclusions

We developed an implantable bioelectronic system for continuous, non-anesthetized electrical stimulation immediately after VML injury. NanoPt-modified electrodes enabled safe, daily therapeutic stimulation and concurrent EMG recording throughout the acute post-injury window. Using this system, we demonstrated that early ES preserves neuromuscular structure and restores functional torque to near-normal levels by week 8 in a rodent model of VML, with coordinated improvements in vascularization, macrophage polarization, and satellite cell activation. These findings establish early electrical stimulation as a promising intervention for promoting endogenous muscle regeneration following catastrophic injury.

The magnitude of functional improvement observed here suggests potential translational implications for treating traumatic muscle loss. Neuromuscular electrical stimulation is already used clinically to prevent disuse atrophy during immobilization and to support rehabilitation following stroke and spinal cord injury.^17, 63^ Adapting such approaches for early post-traumatic intervention, potentially through wearable surface electrode systems or minimally invasive implants, could provide a practical pathway for clinical translation. Future work should optimize stimulation parameters while minimizing energy requirements for portable devices, explore alternative ES dosing regimens, and investigate combinations with regenerative biologics (such as growth factors or cell therapies) or engineered scaffolds. The approach developed here provides an essential tool for such combinatorial studies and for systematically dissecting the mechanisms by which electrical activity shapes muscle regeneration.

## Methods

### Animals and ethical approval

All animal procedures were approved by the Carnegie Mellon University Institutional Animal Care and Use Committee (IACUC protocol #202200000015) and conducted in accordance with the National Institutes of Health Guide for the Care and Use of Laboratory Animals. Male and female Sprague-Dawley rats (Charles River Laboratories, Wilmington, MA) aged 10–20 weeks with an initial body mass of 250–500 g were used for all experiments. Animals were housed under standard conditions with a 12-hour light/dark cycle and provided food and water ad libitum. Rats were observed daily for routine clinical parameters, including appetite, activity, and weight-bearing ability on the operated limb.

### Study design and group allocation

Animals were randomly assigned to one of three experimental groups using a computer-generated randomization sequence: VML with electrical stimulation (VML+ES, n = 28), VML without stimulation (VML+NS, n = 28), or non-defect controls receiving stimulation (ND+ES, n = 8). Male and female rats were distributed across all groups; however, the study was not powered to detect sex-specific differences, and sex was not included as a factor in the primary analyses. Investigators performing torque measurements, EMG analysis, and histological quantification were blinded to treatment group assignment. Blinding was maintained until completion of all data collection and primary statistical analyses.

### Implanted system fabrication

The implantable device consisted of two multistranded PFA-coated 304 stainless steel wires (Cooner Wire, Chatsworth, CA; catalog #AS636) and a Pt-Ir reference wire (A-M Systems Inc., Carlsborg, WA; catalog #778000), each 25 cm in length, soldered to a magnetic connector (Adafruit Industries, New York, NY; catalog #5360). Solder junctions were insulated with medical-grade epoxy (LOCTITE HYSOL M-31CL; Henkel, Düsseldorf, Germany).

### Nanoporous platinum electrodeposition

A 25 mM hexachloroplatinic acid (H_2_PtCl_6_) solution was prepared by dissolving anhydrous H_2_PtCl_6_ powder (Sigma-Aldrich, CAS 26023-84-7) in deionized water. Electrochemical deposition was performed using a three-electrode cell configuration with a Pt mesh counter electrode, Ag/AgCl reference electrode, and a 7 mm length of uninsulated multistranded stainless steel wire as the working electrode. Nanoporous platinum (NanoPt) was deposited via cyclic voltammetry, scanning between −0.3 V and +0.3 V versus Ag/AgCl at 0.12 V/s for 100 cycles.

### Electrochemical characterization

All electrochemical measurements were performed inside a grounded Faraday cage using a Gamry Reference 600+ potentiostat (Gamry Instruments, Warminster, PA). A 7 mm uninsulated multistranded wire (before and/or after NanoPt deposition) served as the working electrode, with Pt mesh and Ag/AgCl electrodes as counter and reference electrodes, respectively.

Electrochemical impedance spectroscopy (EIS) was performed from 100,000 to 1 Hz with DC potential of 0 V and AC amplitude of 10 mV. Cyclic voltammetry (CV) was conducted from −0.7 to +0.7 V for bare stainless steel and −0.6 to +0.8 V for NanoPt-modified electrodes at a scan rate of 50 mV/s. Charge storage capacity (CSC) was calculated as the time integral of cathodic current normalized by electrode geometric area.

Charge injection capacity (CIC) was determined from voltage transients measured during symmetric biphasic current pulses (cathodic-first; 1.0 ms cathodic, 0.1 ms interphase delay, 1.0 ms anodic). Maximum cathodic and anodic potential excursions (Emc and Ema) were defined as electrode potentials at the end of each phase when current returned to zero. Charge densities were calculated as the time integral of injected charge per cathodic phase, normalized by geometric area. Cathodic CIC was determined at −0.8 V (stainless steel) or −0.6 V (NanoPt) versus Ag/AgCl; anodic CIC was determined at +0.8 V for both materials.

Electrode stability was assessed in a tissue-mimicking phantom composed of 15% gelatin and 0.2% NaCl in deionized water, formulated to replicate skeletal muscle conductivity and permeability.^64^ Electrodes were inserted 1 mm apart perpendicular to the phantom surface. Biphasic pulses (1 ms per phase, 5 mA, 20 Hz, 50% duty cycle: 5 s on, 5 s off) were delivered using an AM-2100 isolated pulse stimulator (A-M Systems). EIS was performed on NanoPt-coated electrodes before and after 48 hours of continuous stimulation.

### VML defect creation and electrode implantation

Rats were anesthetized with isoflurane (induction 3–4%, maintenance 1.5–2.5%) and placed on a sterile surgical stage. A 10 × 7 × 3 mm partial-thickness defect was created in the mid-third of the right tibialis anterior (TA) muscle adapted from an established protocol.^2^ The excised tissue represented approximately 20% of total TA mass **(Fig. S1)**, calculated using the equation: TA mass (g) = 0.0017 × body mass (g) − 0.0716.^2^

Following defect creation, a trocar was used to tunnel the magnetic connector subcutaneously from the defect site to an incision between the shoulder blades, where it was secured with back sutures. The working and ground leads were sutured 1 cm apart to the muscle belly under the fascia, positioned distal and proximal to the defect site, respectively. The Pt-Ir reference electrode was sutured to the gastrocnemius muscle approximately 3 cm from the working and counter electrodes. The fascia and skin were closed in layers with absorbable sutures.

### Electrical stimulation protocol

Daily stimulation was delivered during the first week (days 0–7), with additional sessions on days 10 and 14 to extend coverage through the proliferative phase of regeneration.

Neuromuscular tetanic contractions were evoked using biphasic constant-current pulses (1 ms per phase) delivered at 20 Hz with a 50% duty cycle (5 s on, 5 s off) via an AM-2100 isolated pulse stimulator. Stimulation was delivered during 10 sessions (days 0–7, 10, and 14), each lasting one hour. On days 0–6, stimulation was delivered through the implanted electrode system. On days 7, 10, and 14, rats were briefly anesthetized with isoflurane, and stimulation was delivered via percutaneous monopolar needle electrodes (Technomed; catalog #TNM-TE-B50600-001) positioned on either side of the peroneal nerve following torque quantification. During each session, current amplitude was adjusted every 10 minutes to maintain visible contraction of the anterior crural compartment.

### Functional torque measurements

Maximum isometric dorsiflexion torque was measured weekly using a custom-built apparatus consisting of a high-precision force gauge (Torbal FC5; Scientific Industries Inc., Bohemia, NY), a 3D-printed threaded footplate to secure the foot, and a fixed leg holder to minimize movement during contractions. Rats were anesthetized with isoflurane (1.5–2.5%) delivered via nose cone throughout the procedure.

Tetanic contractions were evoked at 200 Hz with a 50% duty cycle and 500 ms duration using monopolar stimulation needle electrodes (Technomed; catalog #TNM-TE-AP2535-335) inserted superficially on either side of the peroneal nerve. Current amplitude (typically 0.5–5 mA) was titrated to identify the level generating maximal dorsiflexion force before activation of antagonistic plantar flexor muscles. Following a five-minute rest period, six successive maximal tetanic contractions were recorded.

Torque was calculated as: Torque = F × R × sin(θ), where F is recorded force, R is lever arm length (rat foot length), and θ is foot angle (maintained at 90°, such that sin(θ) = 1). Torque values were normalized to body mass recorded immediately before each measurement session. Peak torque values were identified using a custom MATLAB script (MathWorks, Natick, MA), and the mean of six contractions was calculated per animal per timepoint. Pre-defect torque values served as individual baselines for normalization. Consistency of raw torque values with published literature was confirmed.^65, 66^

### Spontaneous EMG recording and analysis

Spontaneous EMG activity was recorded during free locomotion on days 1 and 4 post-injury. Following magnetic connector attachment, raw EMG signals were recorded in one-minute intervals using an Intan 512-channel controller paired with a bipolar headstage (Intan Technologies, Los Angeles, CA; catalog #C3313) at a sampling rate of 20 kHz.

EMG signals were processed using custom MATLAB scripts. Raw signals were high-pass filtered (second-order Butterworth, 10 Hz cutoff) to remove motion artifacts and notch-filtered at 60 Hz to eliminate powerline noise. Signals were then rectified and low-pass filtered (second-order Butterworth, 5 Hz cutoff) to generate EMG envelopes. Baseline activity was quantified during stable resting intervals, and activation thresholds were set at mean + 2 × standard deviation. Burst events exceeding threshold were identified using MATLAB’s *findpeaks()* function. Quantified parameters included burst count, duration, amplitude, and cumulative integrated area. Peak detection accuracy was verified through visual inspection of representative traces.

### Histological analysis

At designated timepoints (pre-defect, day 3, weeks 1, 2, and 8), rats were euthanized under deep isoflurane anesthesia. TA and EDL muscles were carefully excised, weighed, and fixed in 4% paraformaldehyde containing 0.2% Triton X-100 for 48 hours. TA samples were sectioned parallel to muscle fiber orientation, embedded in paraffin, and cut into 10 µm-thick sections.

Tissue sections were stained with Masson’s trichrome and scanned at 40× magnification using a Motic EasyScan (Motic, Xiamen, China) slide scanner. Tissue composition was quantified within a standardized 1 × 3 mm region of interest encompassing the defect site, adjacent remodeling tissue, and neighboring healthy muscle using QuPath software (version 0.5.1).^67^ A supervised random forest pixel classifier was trained using 20 representative regions each for myofibers, collagen, and non-myofiber nuclei (maximum 50 trees, depth 10, resolution 3.99 µm/pixel). The classifier was applied to quantify relative area fractions of each tissue compartment.

Myofiber cross-sectional area (CSA) was measured manually in QuPath.^67^ For TA and EDL, n = 4 and n = 2 technical replicates were analyzed per biological replicate, respectively, with ≥200 myofibers indexed per slide.

### Immunohistochemistry

Paraffin-embedded sections were baked at 65°C for 1 hour, deparaffinized, and subjected to antigen retrieval in citric acid buffer (pH 6.0) at 100°C for 20 minutes, followed by cooling on ice. Endogenous peroxidase activity was quenched with 3% hydrogen peroxide for 10 minutes at room temperature. Protease-induced epitope retrieval was performed using 1:10,000 protease (Sigma P5147) at 37°C for 10 minutes.

Sections were blocked for 2 hours at room temperature in buffer containing 5% human serum, 2% donkey serum, 2% goat serum, 5% BSA, 5% fish gelatin (Biotium 22011), 0.05% Triton X-100, and 0.05% Tween-20 in TBS. Primary antibodies against CD163 (mouse monoclonal, 1:3,000; Novus NBP110-40686) and Pax7 (rabbit polyclonal, 1:3,000; Aviva ARP32393_P050) were added to sections, which were incubated overnight at 4°C. Sections were then incubated with biotinylated horse anti-mouse IgG secondary antibody (1:1,000; Vector Laboratories BA-2000) for 30 minutes at room temperature, followed by VECTASTAIN ABC reagent (Vector Laboratories PK-7100) for 30 minutes. Signal was developed with DAB substrate (Vector Laboratories SK-4105) for 60 seconds, and sections were counterstained with Hematoxylin QS (Vector Laboratories) for 5 minutes. Slides were dehydrated and coverslipped with Permount mounting medium (Sigma).

Cell density analysis was performed in QuPath,^67^ with n = 4 technical replicates averaged per biological replicate.

### Blood vessel quantification

Blood vessels were quantified within the standardized region of interest using QuPath.^67^ Vessels were defined as structures with cross-sectional area >100 µm^2^, capturing arterioles and venules while excluding capillaries.^52^

### Statistical analysis

All statistical analyses were performed using GraphPad Prism (version 10.4.1; GraphPad Software, San Diego, CA). Data are presented as mean ± standard deviation unless otherwise indicated. Normality was assessed using the Shapiro-Wilk test.

For longitudinal torque data, two-way repeated-measures ANOVA was used with treatment group and time as factors, followed by post-hoc pairwise comparisons using two-tailed Student’s t-tests with Bonferroni correction for multiple comparisons. One-way ANOVA was used to assess torque changes within the ND+ES group over time.

For histological endpoints (myofiber area, vessel density, CD163^+^ cell density, Pax7^+^ cell density), two-tailed unpaired Student’s t-tests were used to compare VML+ES and VML+NS groups at each timepoint. For muscle mass and CSA comparisons among three groups, one-way ANOVA with post-hoc Tukey’s test was used.

EMG burst ratios were compared using two-tailed paired Student’s t-tests (within-group distal versus proximal comparisons) and unpaired t-tests (between-group comparisons at each timepoint).

A significance threshold of *P <* 0.05 was used for all analyses. Exact p-values are reported for significant comparisons. Sample sizes are indicated in figure legends.

## Acknowledgements

T.C.-K. and B.B acknowledge funding support from Defense Advanced Research Projects Agency under Award AWD00001593 (416052-5). The content of the information does not necessarily reflect the position or the policy of the Government, and no official endorsement should be inferred. They also acknowledge support from Materials Characterization Facility at Carnegie Mellon University supported by grant MCF-677785, and Bertucci Nanotechnology Laboratory at Carnegie Mellon University supported by grant BNL-78657879.

## Author contributions

S.G. and T.C.-K. conceived the project. S.G., A.B., R.J, S.V and J.B. performed experiments. S.G., S.V., and J.B. analyzed data. R.G., A.E., L.W., S.J., and D.R. contributed to device fabrication and characterization. M.K. and B.B. provided expertise in VML modeling and histology. D.C-K, A.F., D.W., B.B., and T.C.-K. supervised the work. S.G. wrote the manuscript with input from all authors.

## Data availability

The data supporting the findings of this study are available from the corresponding authors upon reasonable request.

## Code availability

Custom MATLAB scripts used for EMG analysis and torque quantification are available from the corresponding authors upon reasonable request.

