## Supplementary Figures for "Early electrical stimulation promotes functional recovery after volumetric muscle loss"

**This PDF file includes:**

SI Appendix, Figs. S1 to S10

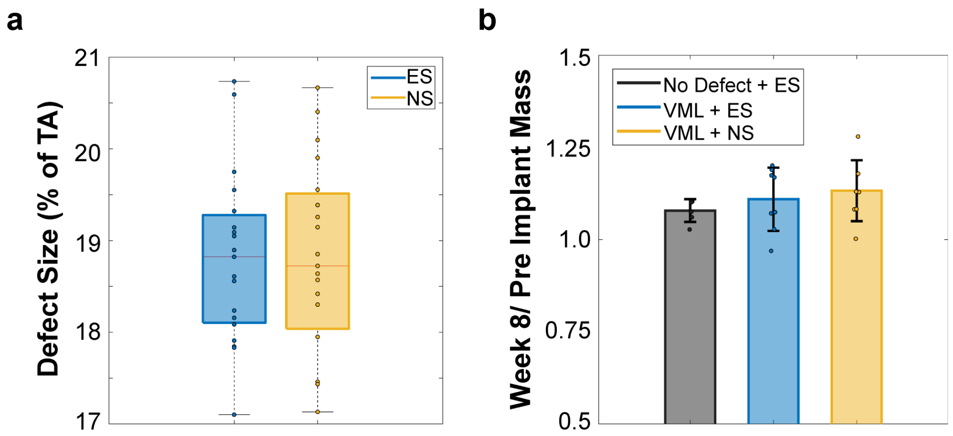


**Fig. S1. Defect mass and body mass at 8 weeks post-implantation**

(**a**) Percentage of TA mass removed for each VML defect (n = 19 per group), calculated as: TA mass (g) = 0.0017 × body mass (g) − 0.0716. (**b**) Ratio of body mass before implantation to 8 weeks post-implantation (n = 6 ND+ES; n = 8 VML+ES and VML+NS). Box plots show median, interquartile range (IQR), and minimum–maximum values. No significant differences were detected (two-tailed Student's *t*-test).

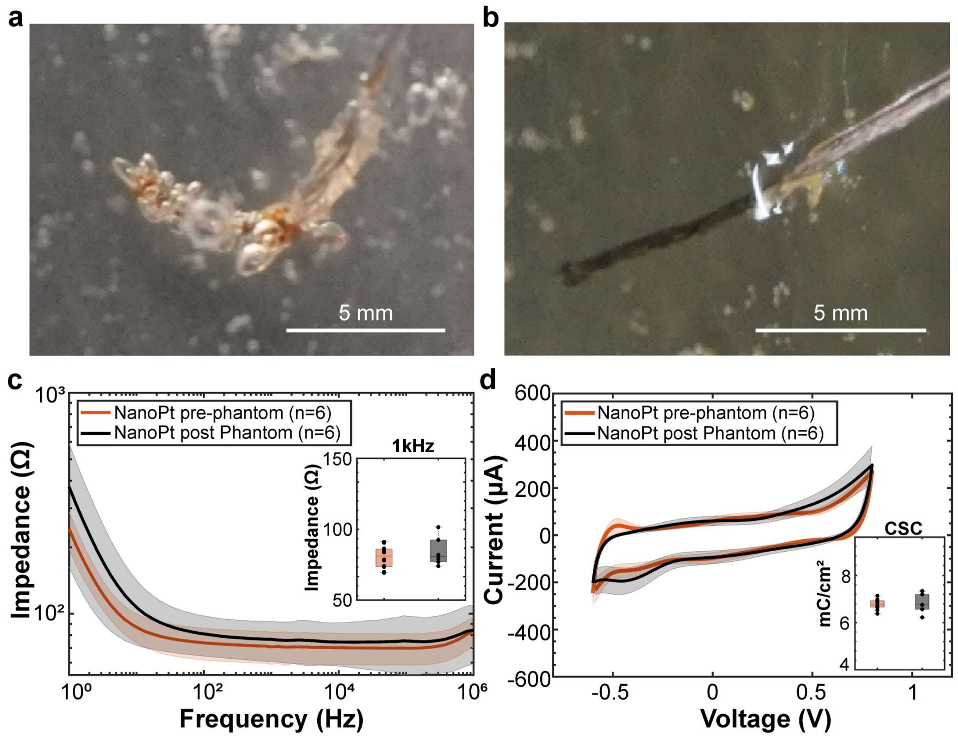


**Fig. S2. NanoPt modification prevents electrochemical degradation during prolonged stimulation in phantom tissue**

Electrochemical characterization was performed in a tissue-mimicking phantom (15% gelatin, 0.2% NaCl) formulated to replicate skeletal muscle conductivity. Biphasic pulses (1 ms per phase, 5 mA, 20 Hz, 50% duty cycle) were delivered via electrodes inserted 1 mm apart. (**a**) Unmodified stainless-steel electrodes exhibited visible gas bubble formation after 3 h of stimulation. (**b**) NanoPt-modified electrodes showed no detectable gas generation or surface oxidation after 48 h of continuous stimulation. (**c**) EIS-derived impedance magnitude and (**d**) cyclic voltammetry confirmed stable electrical properties following prolonged stimulation. Insets show 1 kHz impedance and charge storage capacity (CSC) before and after stimulation. Data are mean (solid line) ± s.d. (shaded region); n values indicated in panels. Box plots show median, IQR, and minimum–maximum values. No significant differences were detected (two-tailed Student's *t*-test).

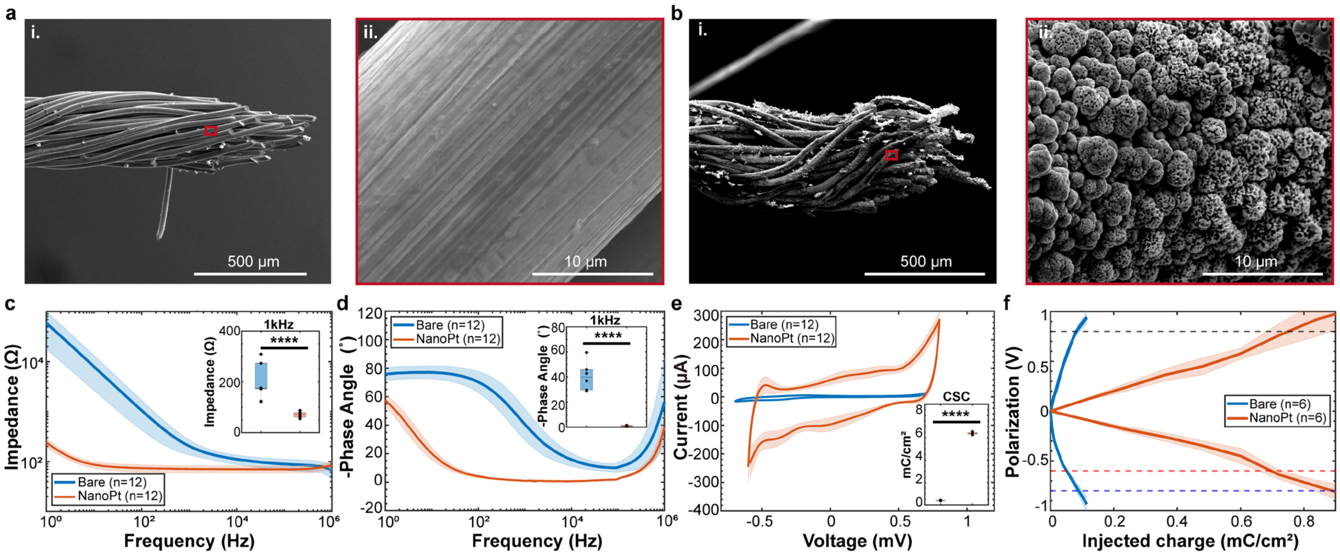


**Fig. S3. NanoPt deposition increases electrode surface area and charge-handling capacity**

Scanning electron micrographs of uninsulated multistrand stainless-steel electrode tips (**a**) before and (**b**) after NanoPt deposition, with (**i**) low and (**ii**) high magnification views. (**c**) EIS impedance magnitude and (**d**) phase angle before and after modification; insets show 1 kHz values. (**e**) Cyclic voltammograms (50 mV $s^{-1}$) with inset showing CSC. (**f**) Charge injection capacity (CIC) comparison. Measurements performed in PBS using Ag/AgCl reference and stainless-steel mesh counter electrode. Data are mean (solid line) ± s.d. (shaded region); n values indicated in panels. Box plots show median, IQR, and minimum–maximum values. $***P < 0.001$, $****P < 0.0001$ (two-tailed Student's *t*-test).

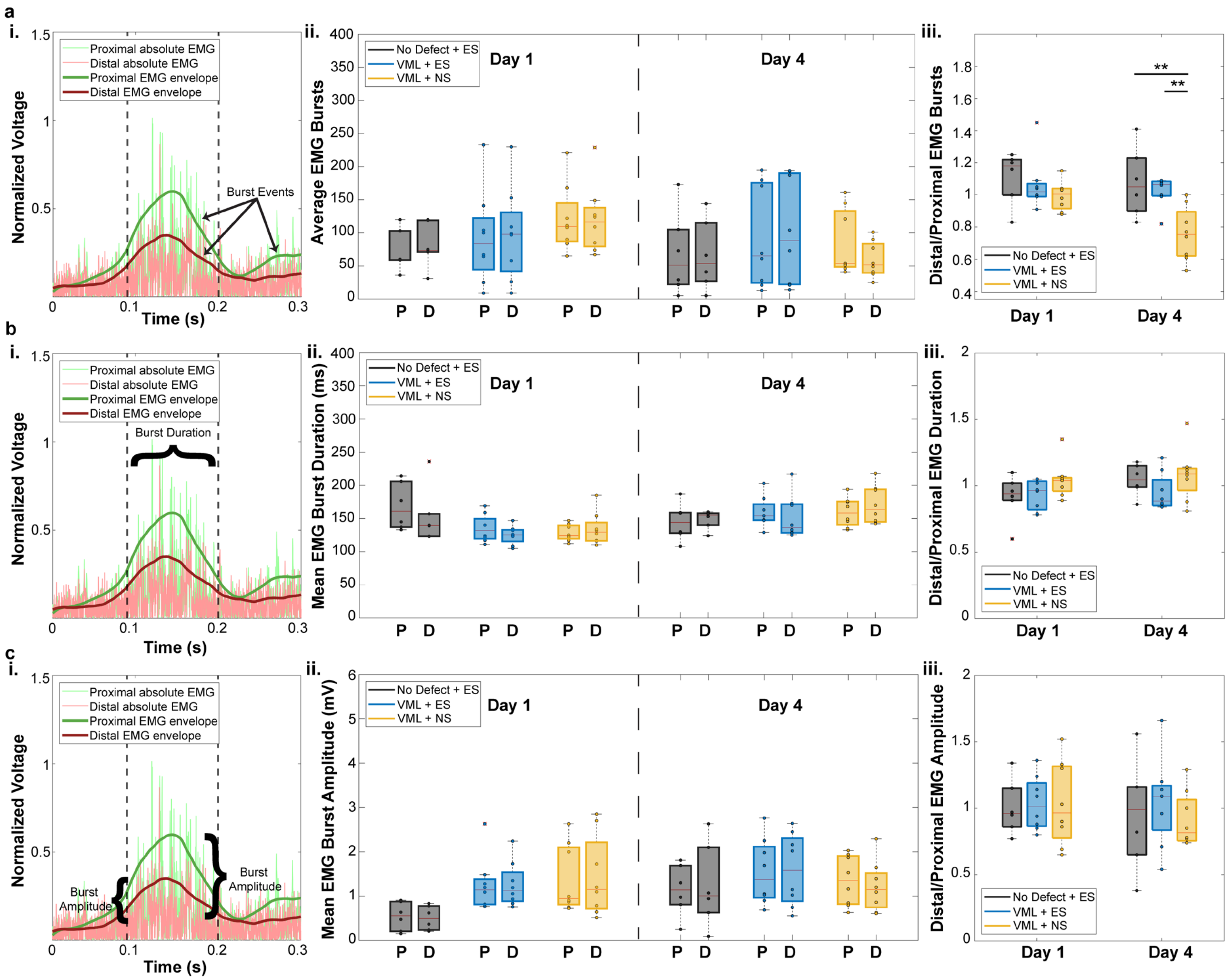


**Fig. S4. EMG burst analysis reveals ES-dependent preservation of distal neuromuscular activity**

Analysis of enveloped EMG recordings showing (**a**) burst events, (**b**) burst duration, and (**c**) burst amplitude. For each parameter: (**i**) schematic of analyzed feature, (**ii**) distributions of proximal (P) and distal (D) values, and (**iii**) distal-to-proximal ratios. Data derived from 1-min recording epochs (n = 6 ND+ES; n = 8 VML+ES and VML+NS). Box plots show median, IQR, and minimum–maximum values; outliers indicated by red "×". $*P < 0.05$, $**P < 0.01$, $***P < 0.001$ (two-tailed Student's $t$-test).

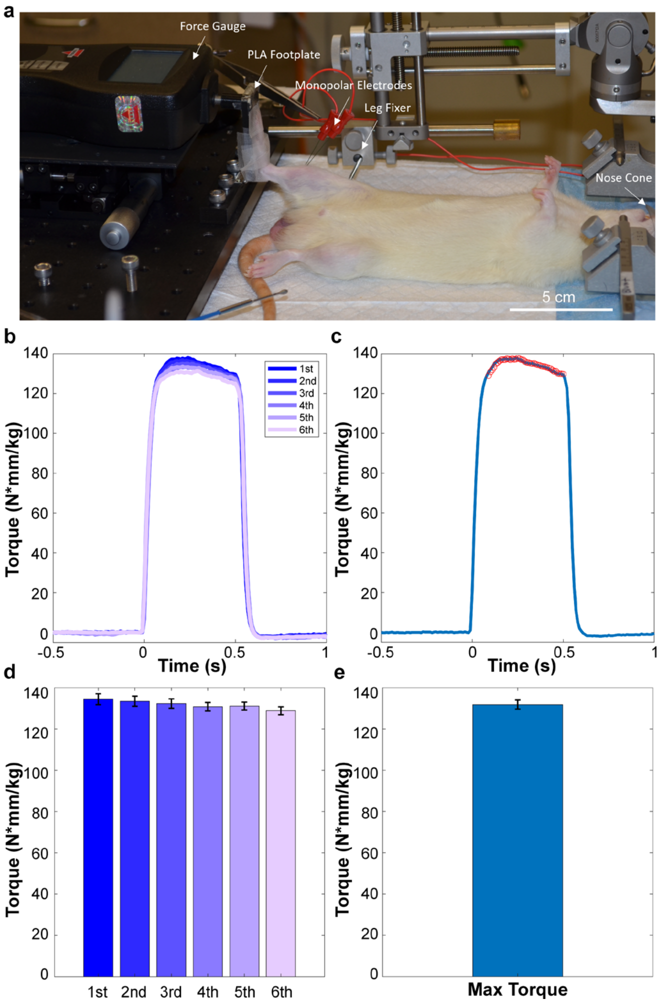


**Fig. S5. Torque measurement system and data acquisition**

(**a**) Custom apparatus consisting of force gauge, 3D-printed footplate, monopolar stimulation electrodes, and fixed leg holder. (**b**) Following current titration to identify maximal response, six successive tetanic contractions were evoked (200 Hz, 500 ms duration) after a 5-min rest period. (**c**) Representative traces showing torque peak identification. (**d**) Individual and (**e**) composite maximal torque values. Bar graphs show mean ± s.d.

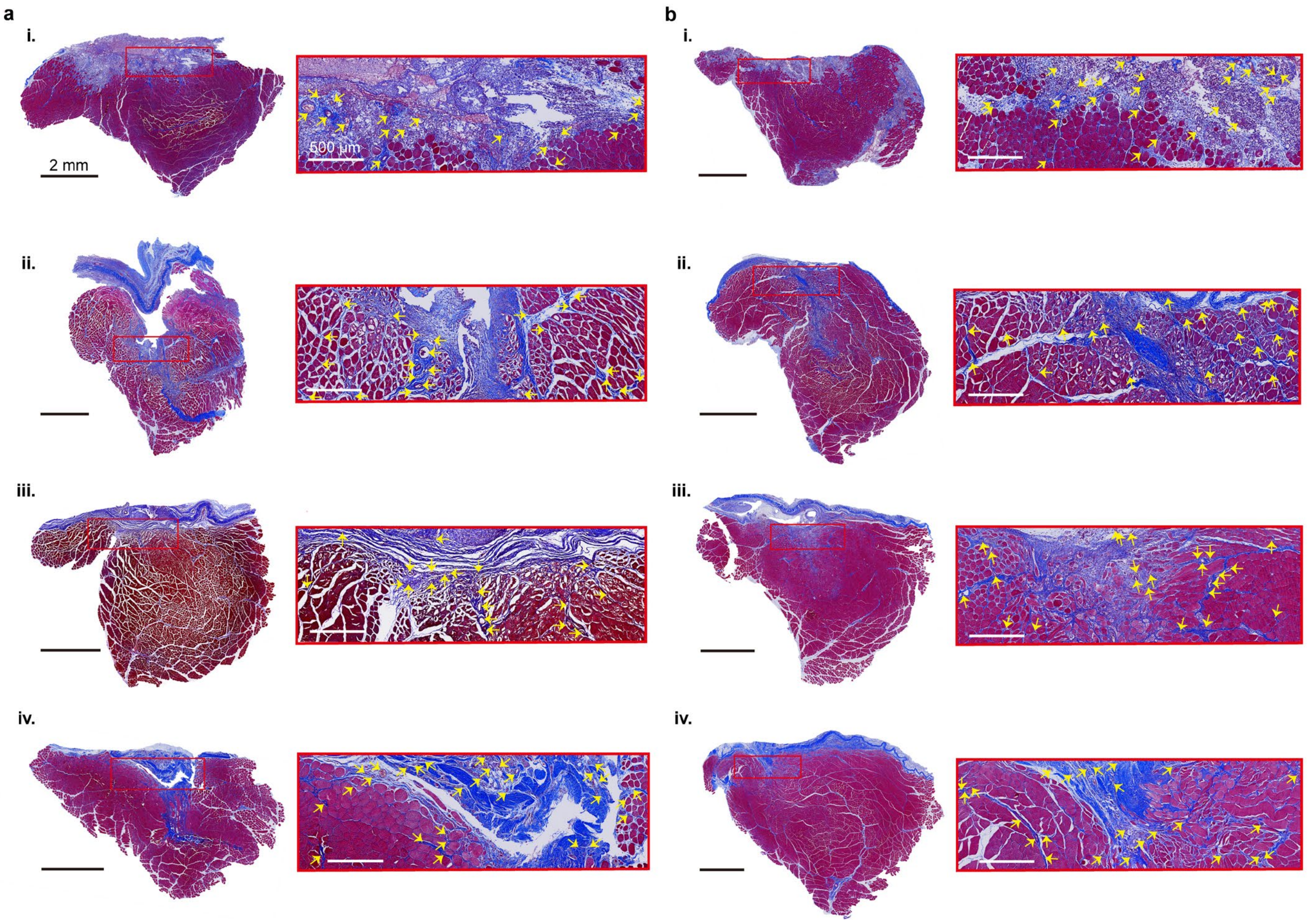


**Fig. S6. Early ES promotes vascular remodeling after VML**

Representative Masson's trichrome cross-sections of TA muscles from (**a**) VML+NS and (**b**) VML+ES groups at (**i**) day 3, (**ii**) week 1, (**iii**) week 2, and (**iv**) week 8. Outlined regions (1 × 3 mm) indicate areas used for compositional and immunohistochemical analysis; higher magnification shown below each panel. Yellow arrows indicate mature and developing blood vessels.

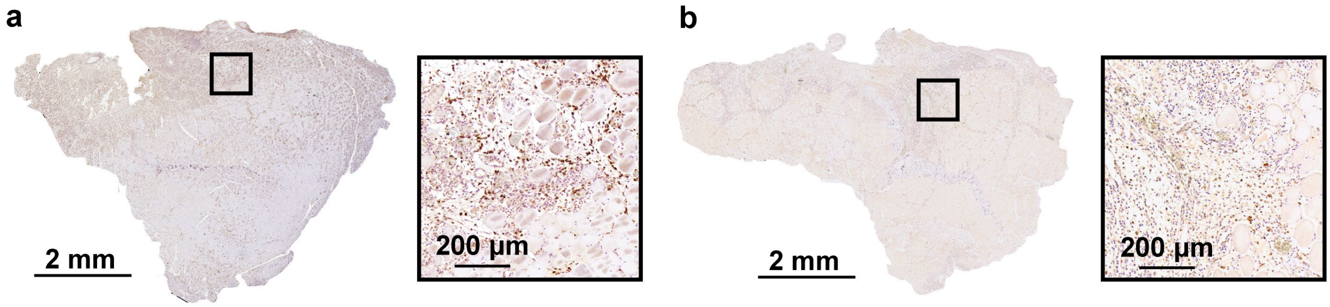


**Fig. S7. CD163 immunohistochemistry at day 3 post-VML**

Representative cross-sections of TA muscle stained for CD163 (M2 macrophage marker) in (**a**) VML+ES and (**b**) VML+NS groups. Insets show magnified views of highlighted regions.

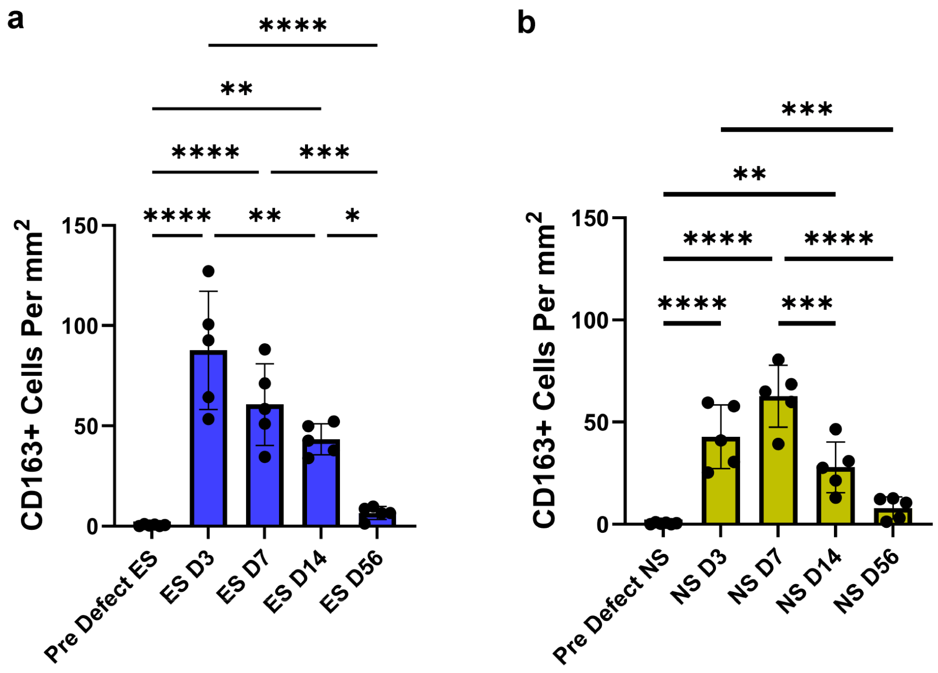


**Fig. S8. Temporal dynamics of $CD163^{+}$ macrophage density**

$CD163^{+}$ cell density in (**a**) VML+ES and (**b**) VML+NS groups at pre-defect, day 3, and weeks 1, 2, and 8. Each point represents n = 5 biological replicates (mean of four technical replicates per animal). *$P < 0.05$, **$P < 0.01$, ***$P < 0.001$, ****$P < 0.0001$ (one-way ANOVA).

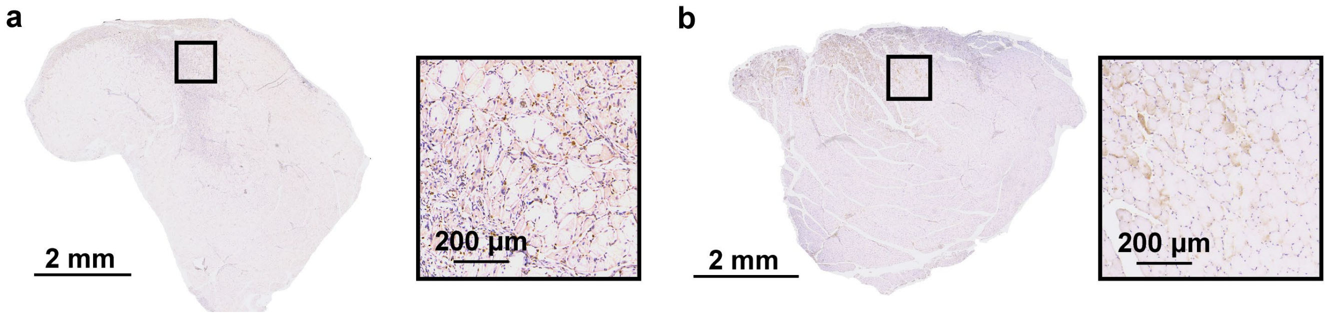


**Fig. S9. Pax7 immunohistochemistry at week 1 post-VML**

Representative cross-sections of TA muscle stained for Pax7 (satellite cell marker) in (**a**) VML+ES and (**b**) VML+NS groups. Insets show magnified views of highlighted regions.

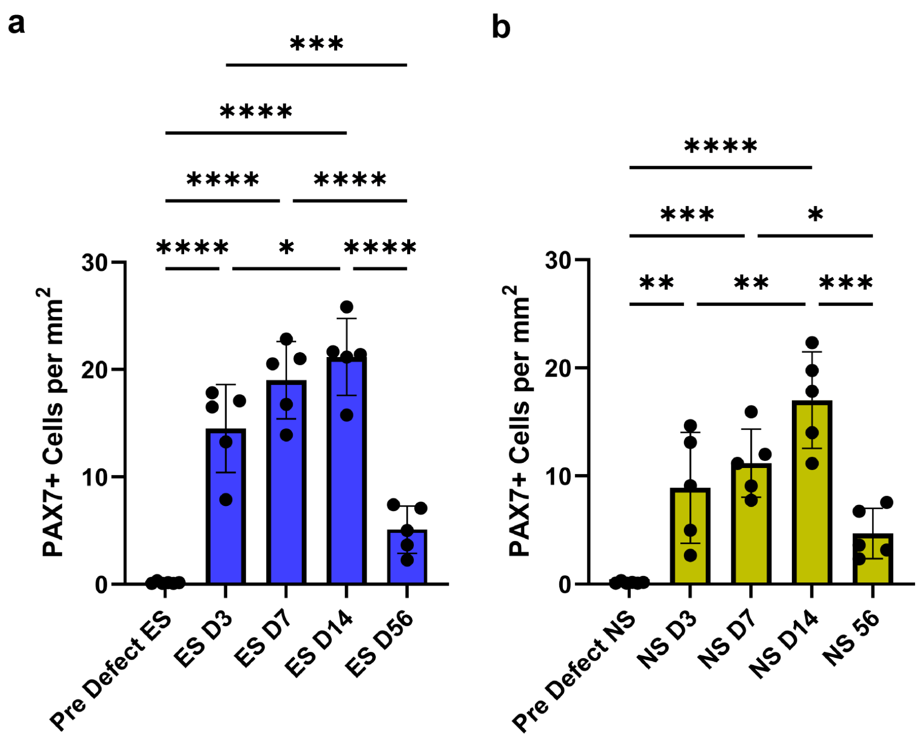


**Fig. S10. Temporal dynamics of Pax7⁺ satellite cell density**

Pax7⁺ cell density in (**a**) VML+ES and (**b**) VML+NS groups at pre-defect, day 3, and weeks 1, 2, and 8. Each point represents n = 5 biological replicates (mean of four technical replicates per animal). **P* < 0.05, ***P* < 0.01, ****P* < 0.001, *****P* < 0.0001 (one-way ANOVA).

**Data S1. (separate file)**

Data and statistical tests for Manuscript panels.

**Data S2. (separate file)**

Data and statistical tests for Supplemental Material panels.
